# Cross-disorder genetic analysis of immune diseases reveals distinct disease groups and associated genes that converge on common pathogenic pathways

**DOI:** 10.1101/2022.10.03.510292

**Authors:** Pietro Demela, Nicola Pirastu, Blagoje Soskic

**Affiliations:** Human Technopole, Viale Rita Levi-Montalcini 1, 20157 Milan

## Abstract

Genome-wide association studies (GWAS) have mapped thousands of susceptibility loci associated with immune-mediated diseases, many of which are shared across multiple diseases. To assess the extent of the genetic sharing across nine immune-mediated diseases we applied genomic structural equation modelling (genomic SEM) to GWAS data. By modelling the genetic covariance between these diseases, we identified three distinct groups: gastrointestinal tract diseases, rheumatic and systemic diseases, and allergic diseases. We identified 92, 103 and 91 genetic loci that predispose to each of these disease groups, with only 12 of them being shared across groups. Although loci associated with each of these disease groups were highly specific, they converged on perturbing the same pathways, primarily T cell activation and cytokine signalling. Finally, to assess whether variants associated with each disease group modulate gene expression in immune cells, we tested for colocalization between loci and single-cell eQTLs derived from peripheral blood mononuclear cells. We identified the causal route by which 47 loci contribute to predisposition to these three disease groups. In addition, given that the assessed variants are pleiotropic, we found evidence for eight of these genes being strong candidates for drug repurposing. Taken together, our data suggest that different constellations of diseases have distinct patterns of genetic association, but that associated loci converge on perturbing different nodes in a common set of T cell activation and signalling pathways.

## Introduction

Immune-mediated diseases are chronic and disabling conditions where the immune system attacks healthy tissue, leading to its destruction. It is well documented that these diseases co-occur within families and that multiple immune diseases are likely to occur in the same individual ^1–3^ suggesting that immune diseases have a shared genetic basis.

Genome-wide association studies (GWAS) have identified thousands of susceptibility loci associated with immune-mediated diseases, many of which have been observed in multiple diseases ^4,5^. For example, the major histocompatibility complex (MHC) locus is associated with most of autoimmune diseases ^6^. Another example is a locus containing *CTLA4* which is associated with multiple immune diseases including rheumatoid arthritis (RA), celiac disease (CeD), type 1 diabetes (T1D) and Hashimoto thyroiditis (Ht) ^7–10^. Targeting the CTLA-4 pathway has been successful in tumour immunotherapy, however in more than 60% of patients, CTLA-4 blockade leads to multiorgan autoimmune reaction ^11^. In contrast, the property of CTLA-4 to bind the costimulatory molecules is extensively used as a treatment for RA ^12^.

Understanding the pleiotropy of genetic associations is critical, as it can reveal common disease mechanisms and pathogenic pathways. A cross-disorder genomic analysis could identify shared mechanisms and potential targets for drug repurposing. By combining cases and controls across immune diseases, recent work identified 224 shared associations, improved fine-mapping and revealed shared disease genes such as *RGS1* ^13^. Similarly, a study using local genetic correlation showed widespread sharing across traits ^14^. For example, T1D and Systemic Lupus Erythematosus (SLE) shared 18 loci. Another study assessed the regulatory activity of immune disease associated SNPs and showed that shared genes were highly connected and were involved in immune pathways ^15^. Although it has been established that immune phenotypes have a shared genetic predisposition, further detailed and systematic analysis is necessary to understand the causes and structure of such sharing. In particular, it is unclear whether sharing is equally distributed across immune diseases (i.e. is there a common factor conferring general risk for all immune disease?) or there are subgroups of immune diseases that are more similar to each other than the rest.

Here we sought to investigate common factors representing general risk across immune diseases. To examine the genetic architecture of nine immune-mediated diseases we applied genomic structural equation modelling (genomic SEM) ^16^ to GWAS data. This revealed three groups of diseases: first consisting of diseases affecting the gastrointestinal tract, the second consisted of rheumatic and systemic disorders and the third group contained allergic diseases.

Each group had an unique genetic architecture and only a handful of loci were in common among the groups. Collectively, our results provide new insights into shared mechanisms of genetic risk for immune-mediated diseases and prioritise drug targets that could be used for multiple disorders.

## Results

### Factor analysis reveals three groups of immune-mediated diseases

To investigate whether there is a common genetic factor underlying multiple immune-mediated diseases, we first used the multivariate LD score regression implementation in genomic SEM ^16,17^ to estimate genetic correlations among nine diseases (Crohn’s disease, CD; ulcerative colitis, UC; primary sclerosing cholangitis, PSC; juvenile idiopathic arthritis, JIA; systemic lupus erythematosus, SLE; rheumatoid arthritis, RA; type 1 diabetes, T1D; eczema, Ecz; asthma, Ast) (Figure 1A, Supplementary Table 1). We collected GWAS summary statistics for each of the traits, and we selected studies that used genome-wide genotyping arrays, as it is required for accurate estimation of LD score regression. We then modelled the genetic variance-covariance matrices across traits using genomic SEM ^16^. This allowed us to uncover latent factors which represent shared variance components across diseases (Figure 1B). By using a range of model fit statistics, we were able to show that the genetic correlation structure was best described by a model using three factors (Supplementary Figure 1A-C). Factor one consisted of diseases affecting the gastrointestinal tract (CD, UC and PSC). Factor two contained autoimmune diseases, which were largely rheumatic and systemic disorders (RA, SLE, JIA and T1D). Finally, factor three contained allergic diseases (Ast and Ecz) (Figure 1B). Therefore, we refer to factors as: F_gut_, F_aid_ and F_alrg_.

**Figure 1.**
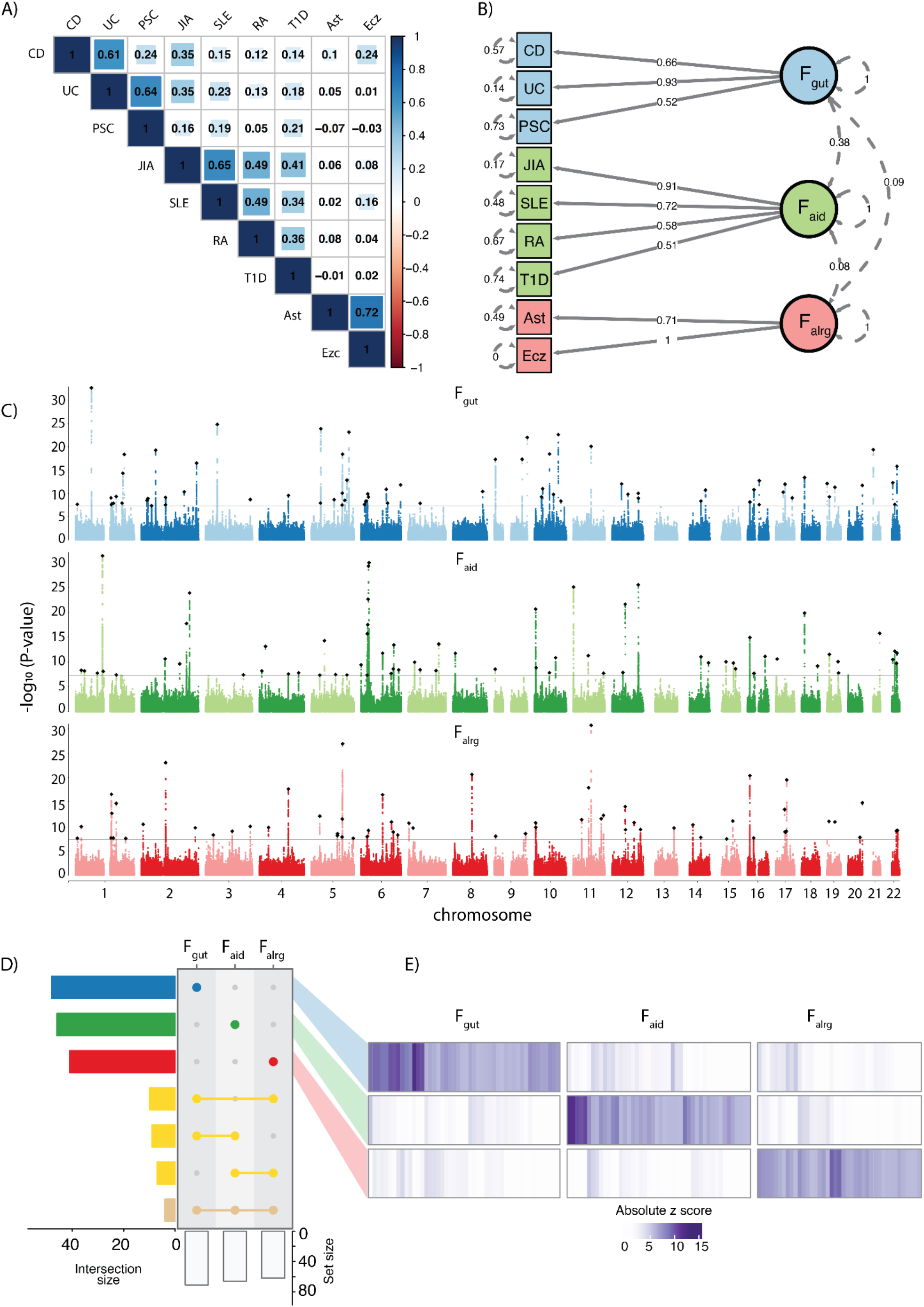
Three groups of immune-mediated diseases have distinct patterns of genetic associations. **A**) Genetic correlation matrix of nine immune-mediated disorders estimated with LD score regression. Shades of blue and red indicate positive and negative correlations respectively. (**B-D**) Blue represents F_gut_, green F_aid_ and red F_alrg_. **B**) Path diagram of the three-factor model of immune-mediated diseases. Colours represent different factors. Latent variables representing common genetic factors are depicted as circles. Standardised loadings (one-headed arrows), residual variances (two-headed arrows connecting the variable with itself) and covariances (two-headed arrows connecting latent variables) are shown. **C**) Manhattan plots of SNP-specific effects on each factor. Black rhomboids represent lead SNPs and a solid line indicates the genome-wide significant threshold (p-value= 5×10^−8^). **D**) UpSet plot showing the overlap between significant genomic regions associated with different factors; intersection size indicates the number of overlapping regions. Asymmetric overlaps (e.g. two regions in one factor overlapping with one region in the other) are counted as one overlap. Yellow represents overlapping genomic regions. **E**) Heatmap of absolute z-scores of factor specific genomic regions. Each column corresponds to a lead SNP, with rows corresponding to factors. Hierarchical clustering was applied to the columns, with breaks along columns separating the factor-specific lead SNPs. CD, Crohn’s disease; UC, ulcerative colitis; PSC, primary sclerosing cholangitis, JIA, juvenile idiopathic arthritis; SLE, systemic lupus erythematosus; RA, rheumatoid arthritis; T1D, type 1 diabetes; Ecz, eczema; Ast, asthma.

To identify how genetic variation impacts the identified latent factors, we tested the association between common SNPs across GWAS studies and each of the latent factors. We discovered 201 genome-wide significant regions that are associated with latent factors, 72 for F_gut_, 66 for F_aid_ and 63 for F_alrg_ (Figure 1C and 1D and Supplementary Table 2). Strikingly, the overlap between these regions was modest, with only 30 out of 201 genomic regions overlapping among at least two factors, and only four regions overlapping across all three factors (Figure 1D). Comparing the z-scores for the three factors within each region showed that this modest overlap was not due to p-value thresholding (i.e the same region in another factor having a p-value just below the threshold) (Figure 1E). In addition, we correlated F_gut_, F_aid_ and F_alrg_ with psoriasis ^18^ and allergies ^19^ GWAS, and showed that they have high correlation with F_alrg_ and not with the other factors (Supplementary Figure 2A). Furthermore, eosinophil counts ^20^ also showed the highest correlation with F_alrg_, giving further support to our factor definition (Supplementary Figure 2B). We did not observe strong genetic correlation with lymphocyte or monocyte counts ^20^ (Supplementary Figure 2B).

Finally, we investigated whether the SNPs were acting via the three factors according to the proposed causal model or, whether SNPs had independent effects on the diseases that the factors are composed of. To do so, we computed the Q_SNP_ heterogeneity statistics (Methods and ^21^). In short, Q_SNP_ allows us to identify SNPs that plausibly do not affect individual diseases exclusively by their associations with the latent common factors. In other words, if the Q_SNP_ heterogeneity statistic is significant, it implies that the tested SNP acts at least partially independently of the latent factors. Our results show that only 10% of loci were significant for Q_SNP_ heterogeneity (22/201) (Supplementary Figure 3A), suggesting that the three factor model explained the genetic structure at the individual SNP level for 90% of identified regions.

### Latent factors have a distinct genetic architecture

An overlap of GWAS regions across two traits does not imply that the underlying causal mechanism is the same across traits. Given that many GWAS regions are complex and could contain multiple independent signals, we performed a systematic analysis of identified regions by combining conditional analysis with colocalization. Briefly, to increase the robustness of colocalization, we devised a statistical approach where the association signal is first decomposed into its conditionally independent components. Next, each component was used for colocalization testing which allowed us to group similar association signals (Figure 2A). This approach enabled resolving complex regions and discovering colocalization events for secondary signals, which would not have been possible by colocalizing the whole regions. Due to challenges of the HLA region we removed two genomic regions encompassing *HLA* genes. We identified 286 independent signals in 199 GWAS associated regions (Supplementary Table 3-6). Out of these 286 loci, 84 were specifically associated with F_gut_, 94 with F_aid_ and 83 with F_alrg_ (Supplementary Table 3-4 and Figure 2B). Only 11 loci were shared across any two factors, and only one was shared across all 3 factors. This further demonstrated that each group of diseases had a specific pattern of genetic associations. For example, a region on chromosome 16 encompassing multiple genes (11,006,011−11,751,015) had significant associations with all three factors (Figure 2C and 2D). However, the conditional analysis and colocalization demonstrated that these signals are independent and not shared across factors. In this region we identified three independent signals that colocalize between CD and F_gut_: rs12922863 (the closest gene *CIITA* which is involved in antigen presentation), rs415595 (the closest gene *TNP2* involved in the regulation of protein processing) and rs13335254 (the closest gene *LITAF* which regulates TNF-alpha expression). Similarly, F_aid_ and F_alrg_ had two independent signals each, which colocalized with T1D and Ecz respectively. The locus that was shared across all three groups of diseases is located at chromosomes 4 (122,903,441-123,720,933) and encompasses a potent regulator of T and B cell proliferation *IL21*.

**Figure 2.**
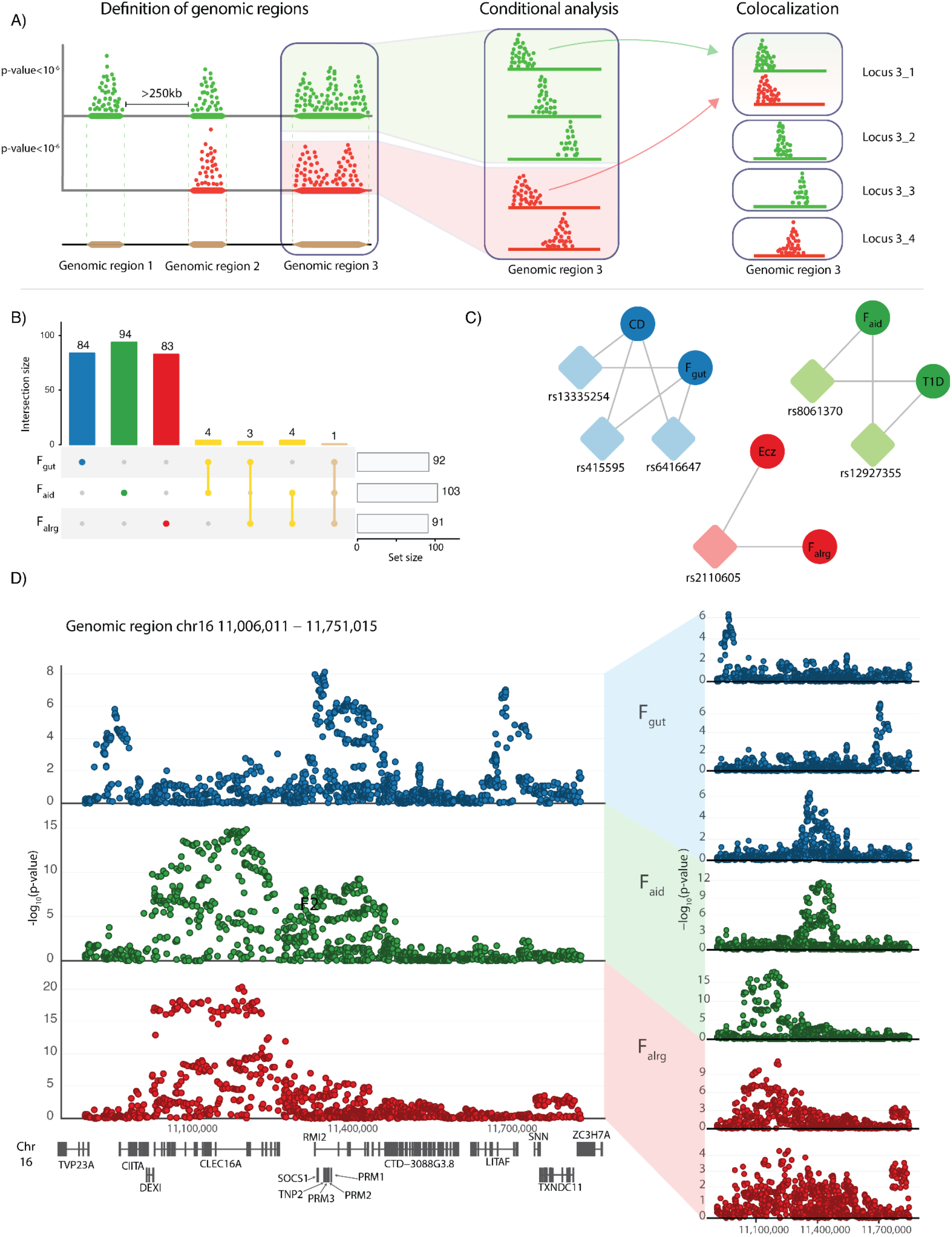
Latent factors have a distinct genetic architecture. **A)** Diagram of the conditional analysis and colocalization strategy (see Methods). Colours represent different traits. **B)** Blue, green and red represent loci that were specific for F_gut_, F_aid_ and F_alrg_ respectively, while yellow represents loci that are shared between factors. **C)** Colocalization relationship between latent factors and traits in the region 16:11,006,011 − 11,751,015. Colours represent disease groups. Circles represent latent factors or traits, rsID of the lead SNP and rhomboids represent the loci that colocalize among traits. **D)** Conditional analysis of the genomic region 16:11,006,011−11,751,015. Locus-zoom plots of three different factors (blue for F_gut_, green for F_aid_, and red for F_alrg_) and the conditional loci for each of the latent factors in the regions are shown.

Taken together, we identified independent signals between factors and determined how each of the factors relate to individual diseases and their likely causal genes.

### Factor-associated loci perturb different nodes in T cell activation and signalling

Identifying transdiagnostic risk pathways can uncover critical cell functions whose perturbations lead to immune system dysfunction and diseases. Therefore, we sought to translate factor-associated variants to cellular functions. Briefly, we identified the closest gene to the lead SNP within each locus and used these genes to test for pathway enrichment with gProfiler2 (Methods). Genes within the associated loci were enriched in cytokine signalling, differentiation of T helper cells, and various immune diseases as well as response to pathogens (Figure 3A and Supplementary Table 7). Given the modest overlap of factor-associated loci, we expected that the enriched pathways would be distinct across factors. However, factor associated genes were largely enriched in the same pathways, although different genes were driving this enrichment (Figure 3A). For example, we observed that both F_gut_ and F_aid_-associated loci were enriched in the JAK-STAT signalling pathway, which is critical for response to many cytokines (Figure 3B). Despite both F_gut_ and F_aid_ being enriched for JAK-STAT signalling, the implicated genes were distinct. Notably, several loci encompassing cytokine genes (*IL2*, *IL10*, *IFNG*, *IL12B*) were associated with the F_gut_ group of diseases, while only *IL21* was associated with the F_aid_ group of diseases. Similarly, transcription factors *STAT1* and *STAT4* were specifically associated with F_aid_, while *STAT3* was associated with F_gut_. This suggests that although trans-diagnostic risk loci are different for three groups of diseases, they converge on perturbing similar cellular functions.

**Figure 3.**
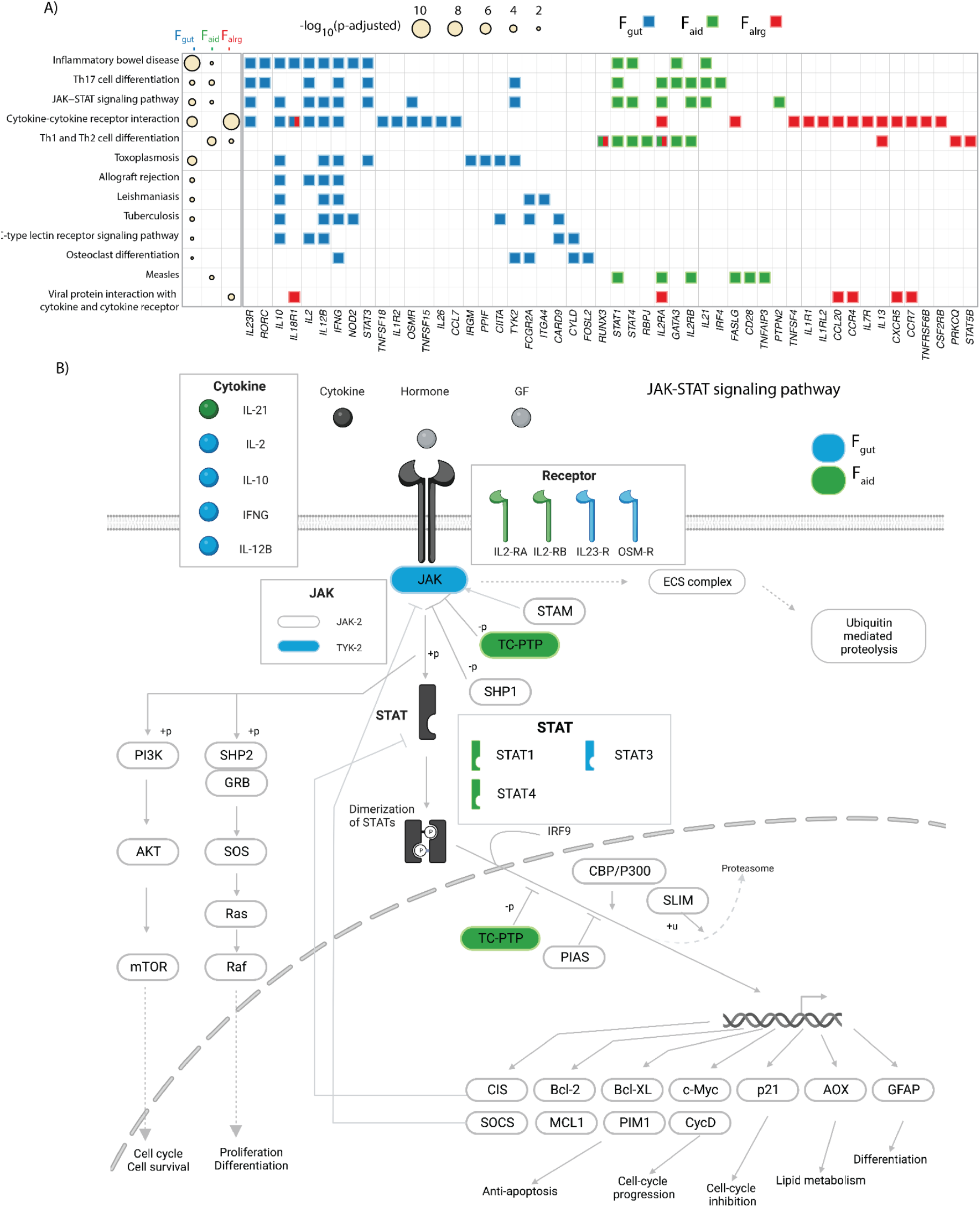
Factor associated loci perturb different nodes of the same pathways. **A**) KEGG pathway enrichment analysis of factor associated genes. The heatmap shows KEGG pathways that were significantly enriched (p-adjusted < 0.05) in factor associated genes. The radius of the circle is proportional to the −log_10_(p-adjusted). The tile plot shows enriched genes in each of the pathways. Blue, green and red represent the genes that contributed to the enrichment for F_gut_, F_aid_ and F_alrg_ respectively. **B)** Schematic representation of JAK-STAT signalling pathway. Blue and green represent components of the pathway that contribute to the enrichment from F_gut_ and F_aid_ respectively.

To test whether transdiagnostic risk variants also converge on a specific cell type, we conducted a MAGMA gene-property analysis implemented in CELLECT ^22,23^. To do that we first used the OneK1K cohort ^24^, which to date is the largest study containing single-cell RNA sequencing (scRNA-seq) data from 982 donors and 1.27 million peripheral blood mononuclear cells (PMBCs). We showed that there is an enrichment of F_gut_, F_aid_, and F_alrg_-associated loci in memory CD4, CD8 and unconventional T cells in all three disease groups (Figure 4A). In contrast, we did not observe enrichment of GWAS loci in naive T cells or B cell populations consistent with previous reports ^25^. Interestingly, NK cells were also enriched, but only for the F_gut_ and F_aid_ group of diseases. A similar pattern of enrichment was observed using S-LDSC (Supplementary Figure 4). In addition, given that tonsils are the secondary lymphoid organs where immune activation occurs, we verified T cell enrichments using a study which profiled human tonsils at the single cell level ^26^. These data showed the same pattern of trans-diagnostic enrichment, observed in CD4 and CD8 T cells (particularly in regulatory T cells) (Figure 4B). As observed in PBMC data, disease loci were generally not enriched in B cells. The exception to that was memory B cells expressing Fc receptor–like-4 (FCRL4+ B cells). FCRL4+ B cells are thought to be tissue resident and have been identified as a potential target in RA therapy ^27^, hence our results provide further genetic support for their modulation. Furthermore, we observed that disease loci were enriched in immune cells from gut ^28^ and lung ^29^ cell atlasses, with the strongest enrichment observed in T cells as previously shown (Supplementary Figure 5A and 5B). Nevertheless, we did not observe enrichment in epithelial or other non-immune cells (Supplementary Figure 5A and 5B). Taken together, the cross disease factors capture true immune signals that are shared across diseases. Finally, we observed a similar enrichment pattern in biological processes across all three groups of diseases. Notably, genes in factor-associated loci were enriched for lymphocyte and immune activation (Figure 4C and Supplementary Table 8), albeit this enrichment was driven by a distinct group of genes (Figure 4D) as demonstrated previously.

**Figure 4.**
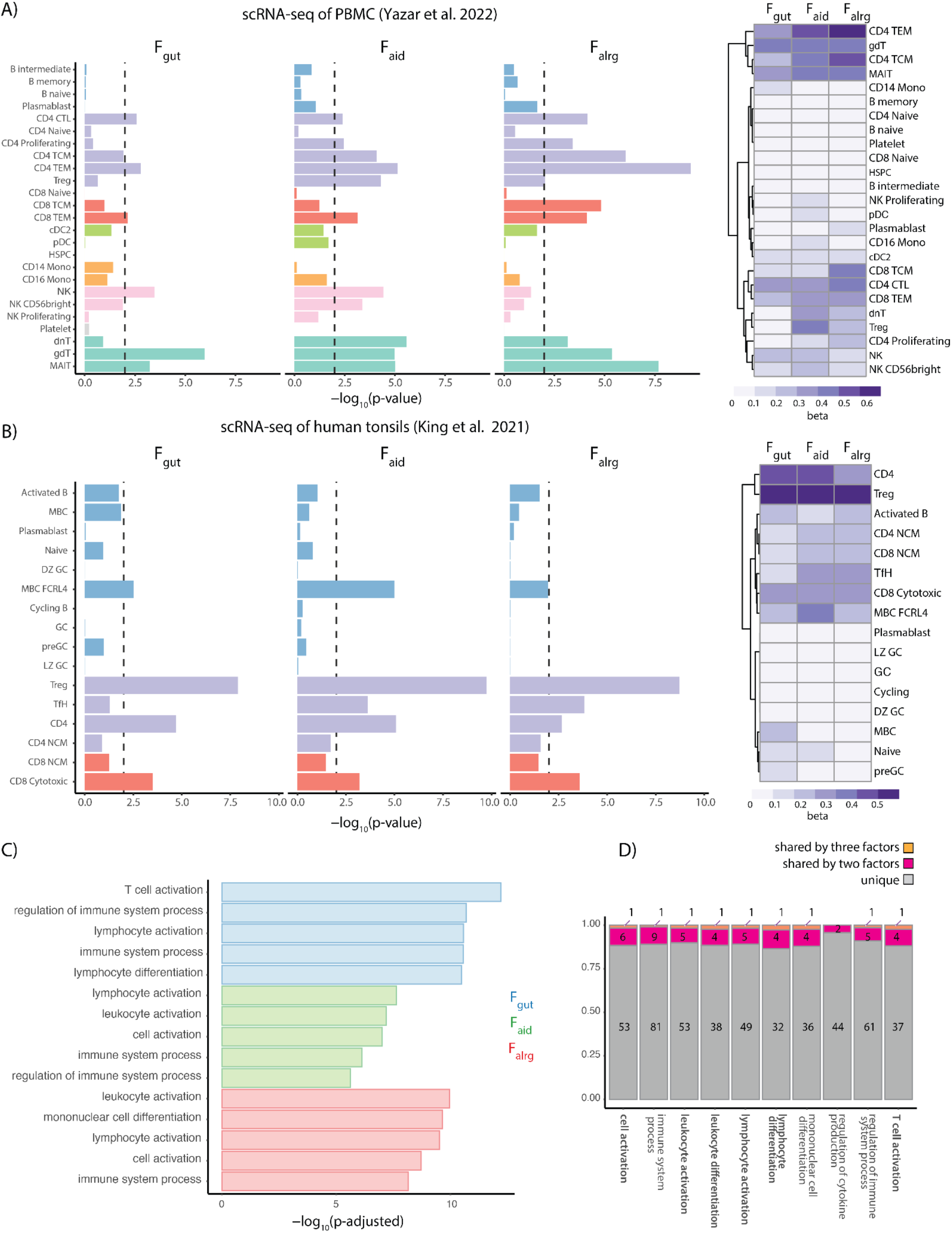
Factor associated loci converge on T cells. **A-B)** MAGMA gene-property results of Onek1k PBMC dataset (**A**) and tonsillar cells (**B**). The barplot shows −log_10_(p-value) of the enrichment. Colours in the barplot represent groups of cells belonging to the same cell-type. The heatmap shows regression coefficients from the MAGMA model. **C)** The bar plot shows the −log_10_(p-adjusted) of the top five GO terms enriched in factor associated genes. Blue, green and red represent the GO terms for F_gut_, F_aid_ and F_alrg_ respectively. **D)** The stacked-bar plot shows the number of genes unique or shared by the latent factors in the top 10 enriched GO terms. We bolded pathways associated with cell activation. Grey represents genes unique to one of the factors, purple represents genes that are associated with two factors and orange represents genes that are associated with all three latent factors.

Taken together, our data suggests that different groups of diseases have distinct patterns of genetic associations but that associated loci converge on perturbing different nodes in lymphocyte activation and cytokine signalling.

### Colocalizing immune cell eQTLs prioritises cross-disease causal genes and identifies potential drug targets

To assess whether variants associated with each disease group modulate gene expression in immune cells, we tested for colocalization between factor-associated loci and single-cell eQTLs (sc-eQTLs) derived from peripheral blood mononuclear cells (PMBCs) from the OneK1K cohort ^24^. Briefly, to identify independent and secondary eQTL signals we performed locus decomposition (see Methods) and colocalized with factor-associated loci using the Bayesian framework *coloc* ^30^. We identified 55 colocalizations in F_gut_, 41 in F_aid_ and 21 in F_alrg_ with PP4 > 0.9 (Supplementary Table 9). Finally, to determine whether an increase of gene expression predicts increased disease risk, we used Mendelian Randomization (MR) using the Wald ratio method (Figure 5A and Supplementary Table 10). For example, an eQTL for Src family tyrosine kinase *BLK* present in naive memory B cells specifically colocalized with an association with the F_aid_ group of traits (Figure 5B), with an increase of *BLK* expression associated with lower disease risk. This is consistent with the fact that rare variants that reduce BLK function have been demonstrated to induce SLE ^31^. In another example, we observed that a locus associated with F_gut_ modulates the expression of Prostaglandin E Receptor 4 *PTGER4* (Figure 5C). In this case, an increase in gene expression is protective to the F_gut_ group of diseases.

**Figure 5.**
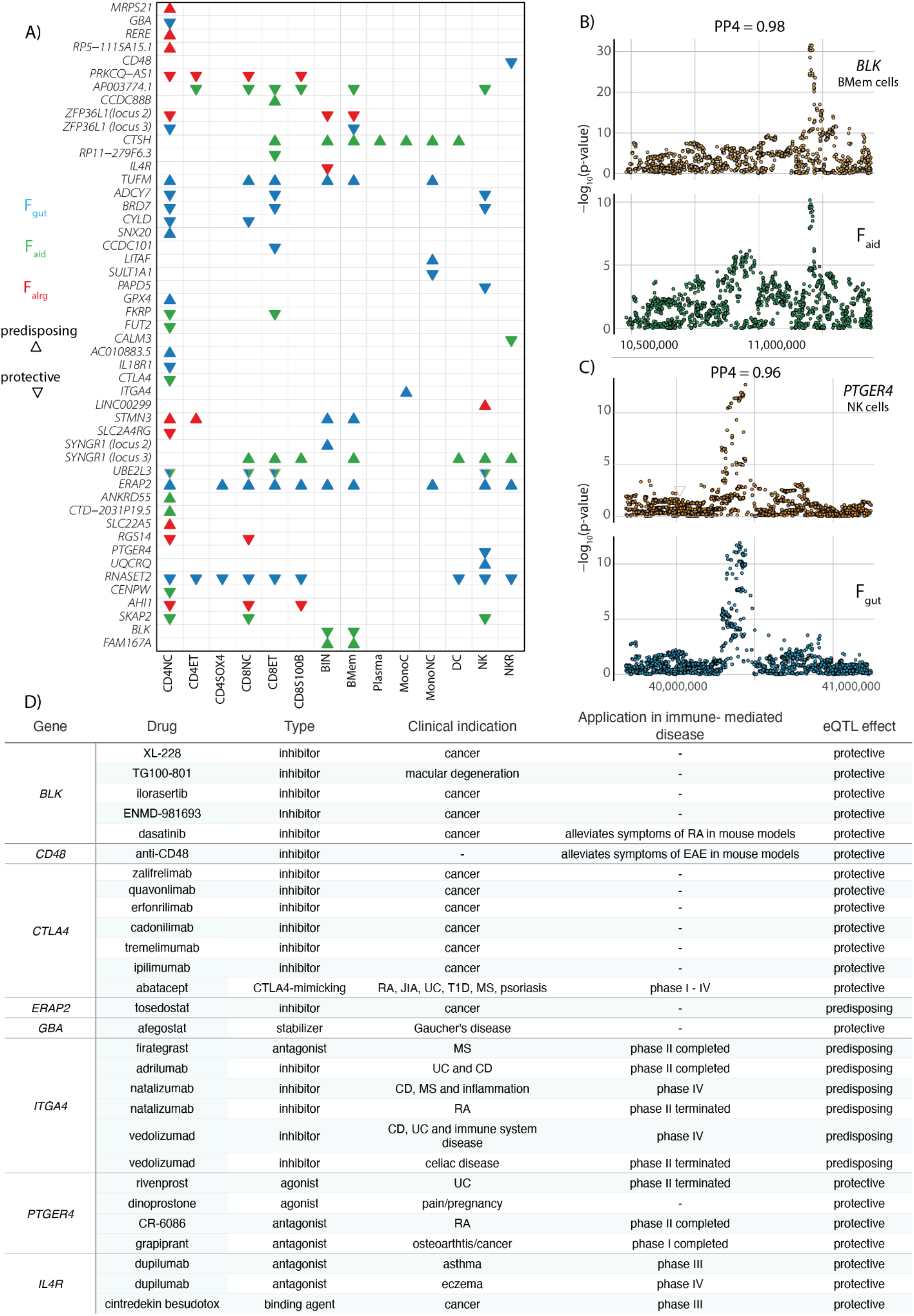
Colocalization of immune cell eQTLs prioritises cross-disease causal genes and identifies potential drug targets. **A)** Colocalization and Mendelian Randomization results (see Methods) of eQTL predicting risk to the latent factors. Triangles pointing upwards indicate that an increase of gene expression increases disease risk, while triangles point downwards indicate decrease of disease risk. Blue, green and red represent F_gut_, F_aid_ and F_alrg_ respectively. Only significant Mendelian Randomization results (p-value <0.05) are shown. **B-C)** Colocalization-plots of latent factors and eQTLs. Posterior probability of colocalization (H4) is shown. **B)** Locus-zoom plot representing the colocalization between the *BLK* gene in B memory cells and F_aid_. **C)** Locus-zoom plot representing the colocalization between the *PTGER4* gene in NK cells and F_gut_. **D**) Table representing the drugs prescribed in clinics, in clinical trials or with preliminary results in mice for immune-mediated disorders targeting eQTL genes. MS, multiple sclerosis; UC, ulcerative colitis; CD, Crohn’s disease; RA, rheumatoid arthritis; JIA, juvenile idiopathic arthritis; T1D, type 1 diabetes; EAE, experimental autoimmune encephalomyelitis.

One of the major hurdles of human genetics has been to translate genetic findings into clinical insights. To identify potential drug targets, we used the Open Targets Platform ^32^ and investigated whether colocalizing genes are known drug targets (Figure 5D). Of the 47 eQTL genes, eight are targeted by drugs which are either already used in the clinics or are in clinical trials. Four of these eight have been previously used in autoimmune diseases, while the other four represent potential candidates for drug repurposing. For example, our data shows that the increase of expression of a key immune regulator *CTLA4* is protective against F_aid_ group of diseases. The property of CTLA-4 to regulate the immune system has long been exploited in treatment of RA ^12^. Similarly, an inhibitor for Integrin Subunit Alpha 4 *ITGA4* has been trailed in UC and CD (Open Targets database and Figure 5D). Our data gives further genetic evidence that increase of *ITGA4* expression leads to an increased risk for F_gut_ diseases, and therefore it is plausible that inhibiting *ITGA4* would be beneficial not only in CD and UC but should also be trialled in PSC.

Taken together, our data shows that understanding the pleiotropy of genetic associations can reveal common disease mechanisms, identify novel drug targets and offer evidence for drug repurposing.

## Discussion

In this work we used genomic SEM to investigate the common genetic factors predisposing to multiple immune-mediated diseases. We identified three broad categories of immune mediated diseases: affecting the gastrointestinal tract, rheumatic and systemic disorders, and allergic diseases. Surprisingly, underlying factors affecting the pathogenesis of each of these disease groups had a highly specific pattern of genetic associations, with only 12/286 loci being shared across these groups. This suggests that there is a genetic similarity between diseases within a group, but that the associated loci are highly distinct across groups. The identified groups agree with previous epidemiological findings. For example, T1D was grouped with rheumatic diseases including RA, which is in line with reports that patients with T1D but not T2D have increased risk of RA (OR=4.9) ^33^. Similarly, approximately 70% of patients with PSC have IBD, with UC being the most prevalent ^34^. Our study shows that there are common genetic mechanisms driving the pathogenesis of these diseases and suggests that creating cross-disorder cohorts of immune diseases could increase the power to identify causal pathogenic processes.

Importantly, over 90% of identified loci acted via common factors, rather than independently on each of the diseases. Therefore, we sought to identify transdiagnostic risk pathways in order to uncover biological processes whose perturbation affects each of the disease groups. Our study showed that despite associated loci being highly factor specific, they converged on perturbing the same pathways involved in T cell activation, differentiation and cytokine signalling. F_gut_ and F_aid_-associated loci were enriched in the JAK-STAT signalling pathway, although there was no overlap in genes driving the pathway enrichment in each of these groups. Loci encompassing cytokine genes (*IL2*, *IL10*, *IFNG*, *IL12B*) and STAT genes (*STAT1* and *STAT4*) were associated with the F_gut_ group of diseases, while *IL21* and *STAT3* were associated with the F_aid_ group of diseases. Similarly, out of 55 genes that are enriched for lymphocyte activation, only 6 were shared across at least two factors. Therefore, one can speculate that perturbations at different nodes which regulate T cell activation and cytokine signalling are partially responsible for driving different disease outcomes. Recent advances in CRISPR editing in T cells and its subpopulations ^35,36^ will be instrumental to elucidate the differential effects of perturbing each node within shared pathways.

Finally, it has been widely demonstrated that supporting preclinical data with genetic evidence can significantly increase the chance of developing successful drugs ^37^. Therefore, understanding how trans-diagnostic variants regulate gene expression can help to identify novel drug targets or supporting evidence to existing trials. Here we colocalized the factor-associated loci with sc-eQTL derived from the OneK1K cohort. To date, OneK1K is the largest study containing single-cell RNA sequencing (scRNA-seq) data from 982 donors and 1.27 million PMBCs. We showed that eight of these colocalizing genes are known drug targets offering further genetic support for their potential therapeutic effect. In addition, given that the assessed variants are pleiotropic, our results imply that identified drugs could be repurposed for diseases within the same group. For example, our data shows that the increase of expression of a key immune regulator *CTLA4* is protective against F_aid_ group of diseases. The property of CTLA-4 to regulate the immune system has long been exploited in treatment of RA ^12^. Similarly, an inhibitor for Integrin Subunit Alpha 4, ITGA4 has been trailed in UC and CD (Open Targets database). Our data gives further genetic evidence that increase of *ITGA4* expression leads to an increased risk for F_gut_ diseases, and therefore it is plausible that inhibiting *ITGA4* would be beneficial not only in CD and UC but should also be trialled in PSC. However, one limitation of this study is that we identified colocalization events for 40 out of 286 loci. This highlights the urgent need for larger cohorts, which will be more powered to detect eQTLs, as well as large-scale genetic studies in immune disease patients.

In conclusion, our work underscores that three groups of immune-mediated diseases do not share similarities in their genetic predisposition, but show associated loci which converge on perturbing different nodes of a common set of pathways, including in lymphocyte activation and cytokine signalling.

## Authors contributions

NP and BS conceived and designed the project. PD, NP and BS performed the data analysis and interpreted the results. NP and BS supervised the analysis. PD, NP and BS wrote the manuscript.

## Acknowledgements

PD is a PhD student within the European School of Molecular Medicine (SEMM). We thank Craig Glastonbury, Cecilia Domínguez Conde, Eddie Cano-Gamez, Laura Esposito, Gosia Trynka and Nicole Soranzo for critical feedback on the manuscript. We also thank Davide Bolognini and Edoardo Giacopuzzi for the computational support.

## Methods

### Processing of summary statistics for LD score regression

We downloaded GWAS summary statistics from published studies on the most common autoimmune disorders: T1D ^7^, RA ^8^, JIA ^38^, SLE ^39^, CD ^40^, UC ^40^, AST ^41^, ECZ ^42^, PSC ^43^ (Supplementary Table 1). Where necessary, rsIDs were added to the summary statistics using the reference file provided in the Genomic SEM repository (https://utexas.app.box.com/s/vkd36n197m8klbaio3yzoxsee6sxo11v/file/576598996073).

Where necessary, chromosomes X and Y were removed and standard error of logistic betas were calculated based on Odds Ratio confidence intervals. Summary statistics were formatted with the *munge* function from Genomic SEM R package v.0.0.5, (with default parameters) which removes all the SNPs not present in the reference file, filters out SNP with MAF < 1% and flips the alleles according to the reference file and computes z-scores. The HapMap3 reference file is provided in the Genomic SEM repository https://utexas.app.box.com/s/vkd36n197m8klbaio3yzoxsee6sxo11v/file/805005013708.

### Estimation of genetic correlation with Genomic SEM

The sum of effective sample size for GWAS that were meta-analysed was calculated by retrieving the information about the cohorts from the respective publications (Supplementary Table 1). We calculated the sample prevalence for each of the cohorts using the following formula

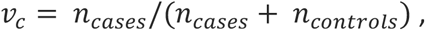

Next, we calculated the cohort specific sample size as follows:

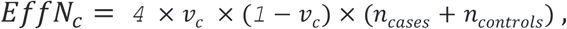

Finally, we summed the *EffN*_*c*_ of each contributing cohort to compute the sum of effective sample size:

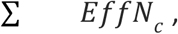

Where *c* are contributing cohorts (as described at https://github.com/GenomicSEM/GenomicSEM) ^44^. To estimate genetic correlation we used the *ldsc* function in Genomic SEM, using the LD reference panel provided in the Genomic SEM repository (https://utexas.app.box.com/s/vkd36n197m8klbaio3yzoxsee6sxo11v/folder/119413852418).

### Factor model specification and GWAS estimation with Genomic SEM

We computed three confirmatory factor analysis models guided by exploratory factor analysis: a) a common factor model with the latent factor variance fixed to 1. b) a two-factor model, where one factor was loading into CD, UC, PSC, JIA, SLE, RA and T1D while the other factor was loading into Ecz and Ast. We allowed for correlation between factors. c) A three factor model where F_gut_ was loading into CD, UC, PSC; F_aid_ was loading into T1D, SLE, JIA, RA, and F_alrg_ loading into Ecz and Ast; we fixed the variance of the latent factors to 1 and allowed correlation between the latent factors (Supplementary Figure 1).

The fit of the model was assessed by estimating the comparative fit index (CFI) and the standardised root mean square residual (SRMR) parameters. We used CFI >0.95 and SRMR < 0.07 as a measure of good fit. Before estimating the SNP-specific effect, we aligned the summary statistics to the reference file (https://utexas.app.box.com/s/vkd36n197m8klbaio3yzoxsee6sxo11v/file/576598996073) which is used to standardise the effect sizes and SE and format the summary statistics (i.e. remove SNPs not present in the reference files and flip the alleles to match the reference) with the *sumstats* function in Genomic SEM with default parameters. SNP-specific effects of 3,309,805 SNPs were estimated with the *userGWAS* function with default parameters using the weighted least squares (WLS) estimation method. In order to evaluate whether the calculated SNP effects were acting through our three factor model, we performed the Q_SNP_ heterogeneity tests. The heterogeneity test returns a *χ*^*2*^, whose null hypothesis suggests that the SNP is acting through the specified model. Therefore, rejecting the null hypothesis means that the SNP acts through a model that is different from the specified one ^16,21^.

### Loci definitions and conditional analysis

We define the boundaries of each significant genomic region by identifying all the SNPs with a p-value lower than 1×10^−6^. We calculated the distance among each consecutive SNPs below this threshold in the same chromosome; if two SNPs were further than 250 kb apart, then they were defined as belonging to two different genomic regions. We then considered as ‘significant’ all the genomic regions where at least one SNP had a p-value < 5×10^−8^. This procedure was repeated for all GWAS. Finally, we compared genomic regions between different GWAS and merged those which overlapped, redefining the boundaries as the minimum and maximum genomic position across all overlapping genomic regions.

### Processing of summary statistics for conditional analysis and colocalization

Before running conditional analysis and colocalization, summary statistics (traits and factors) were processed with the Bioconductor MungeSumstats package ^45^. We specify the parameters to the MungeSumstat function to: align the summary statistics to reference genome to the build GRCh7 (1000genomes Phase2 Reference Genome Sequence hs37d5, based on NCBI GRCh37, R package ‘BSgenome.Hsapiens.1000genomes.hs37d5’ v0.99.1), flip the alleles according to the reference file, remove the SNPs not in the reference file (SNP locations for Homo sapiens, dbSNP Build 144, based on GRCh37.p13, R package ‘SNPlocs.Hsapiens.dbSNP144.GRCh37’ v.0.99.20), exclude the SNPs with betas or standard errors equal to 0.

### Conditional analysis and colocalization

The genomic regions defined in the previous steps are based on genomic position, but multiple association signals may be present within each genomic region. To this end, we developed a statistical approach which first divides each GWAS-significant genomic region into its component signals and then uses colocalization across different traits to group similar association signals. First, in each genomic region for each GWAS we performed stepwise forward conditional regression using COJO ^46^. The stopping criteria was that all conditional p-values were larger than 1×10^−4^. This led to a set of independent SNPs using all SNPs within the genomic region boundary (+/− 100kb). For each SNP, a conditional dataset was produced where SNPs in the genomic region were conditioned to all identified independent SNPs apart from the target one. We then considered as true signals those with p-value p<10^−6^ or those for which the SNP with the lowest p-value was lower than 5×10^−8^ in the original GWAS. This procedure was repeated on all the traits which had a significant association in the considered genomic region. We thus obtained for each trait a set of conditional datasets covering all the SNPs in the genomic region. This procedure is similar to that used by Robinson et al ^47^ but instead of using the step-wise conditioned datasets it uses an ‘all but one’ approach.

To understand which loci were pleiotropic between traits, we ran colocalization using coloc ^30^ analysis between all pairs of loci specific for each trait. Loci which colocalized with PP4 > 0.9 were grouped in a single locus. We excluded the genomic regions in the HLA locus (chromosome 6 - 29,000,000-33,000,000) from this analysis.

### Colocalization with eQTL data

We downloaded eQTLs from the OneK1K cohort ^24^. We first identified for each genomic region if significant cis-eQTLs were present. For each identified eQTL we performed the decomposition of the locus as described above and the identified loci were colocalized with factor and individual trait associated GWAS signals. To claim a true colocalizing signal we required that PP4 > 0.9. In order to identify the direction of the effect of the increase in gene expression for the colocalizing loci, we used Mendelian Randomization using the Wald ratio method (TwoSampleMR R package, ^48^) using as instrument the SNP with the smallest p-value in the conditional datasets. Significant MR results (p-value lower than 0.05) were reported. This procedure was performed cell type per cell type.

### Cell type enrichment

To identify cell types underlying identified factors we used CELL-type Expression-specific integration for Complex Traits (CELLECT). CELLECT quantifies the association between GWAS signal and gene expression specificity using well established models for GWAS enrichment MAGMA ^22^ and S-LDSC ^49^.

### Gene based enrichment

Candidate genes were defined by mapping each lead SNP to the nearest transcription starting site of protein coding genes using the EnsDb.Hsapiens.v75 R package (v2.99.0). To identify enrichment in KEGG pathways, GO terms and REACT pathways we used the R package gprofiler2 (v0.2.1) ^50^, with default parameters. Pathway was considered significant if p-adj < 0.05. We used the R package pathview (v1.34.0) ^51^ to represent the KEGG pathways and to highlight factor-specific genes. The diagram shown in Figure 3B was created with biorender.com using the KEGG pathway as reference.

### Identification of drug targets

Open Targets Platform ^32^ (v.22.06) was used to identify drug targets for eQTL genes. This website was queried on (29th August 2022).

## Supplementary Figures

**Supplementary Figure 1.**
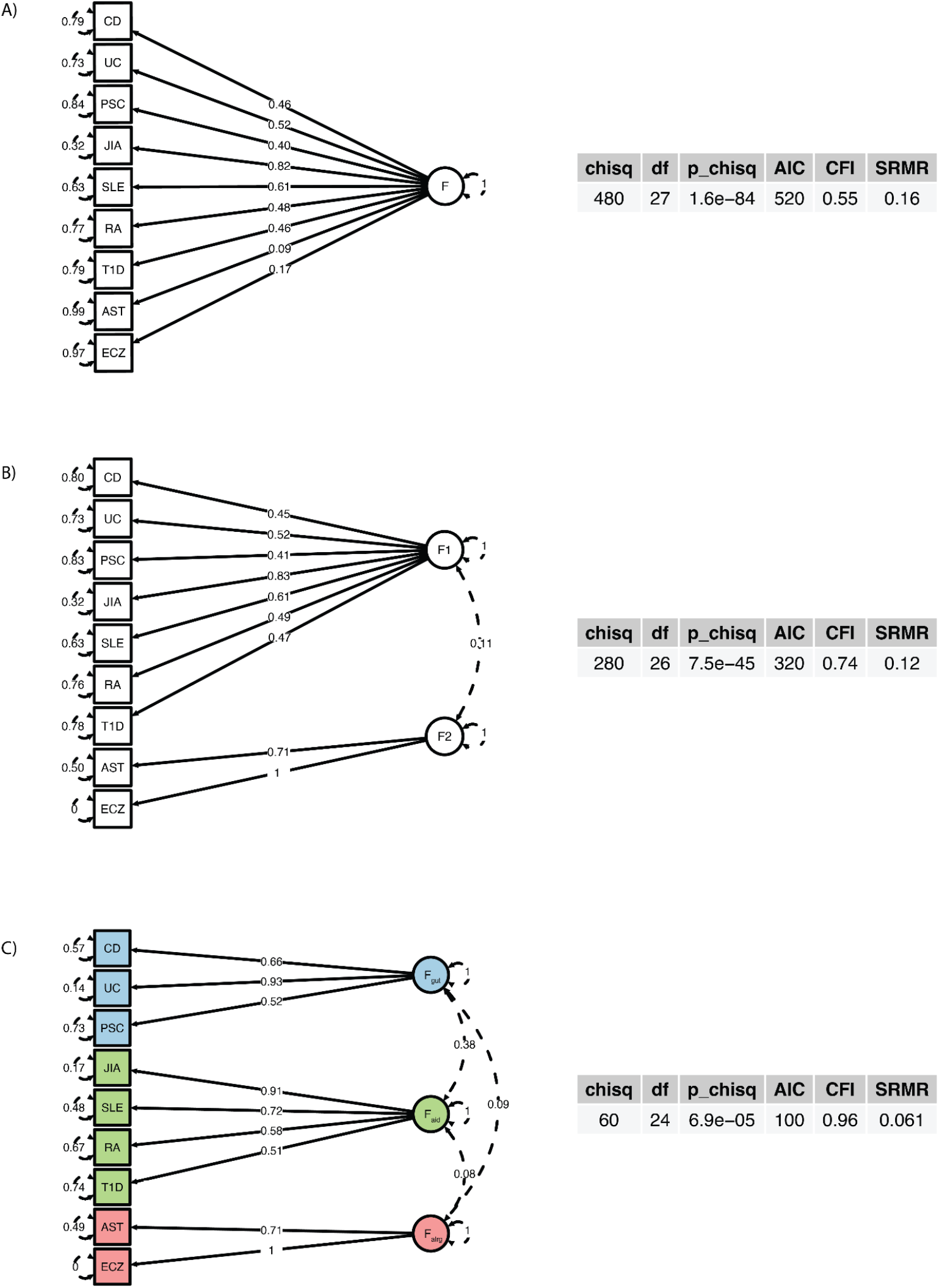
Factor models that were tested and their fit statistics. **A)** A common factor model with the latent factor variance fixed to 1. **B)** a two factor model, where one factor was loading into CD, UC, PSC, JIA, SLE, RA, T1D while the other factor was loading into Ecz and Ast. We allowed correlation between factors and imposed the residual variance to be positive for Ezc. **C**) A three factor model where Fgut was loading into CD, UC, PSC; F_aid_ was loading into T1D, SLE, JIA, RA and F_alrg_ loading into Ecz and Ast; we fixed the variance of the latent factors to 1 and we allowed correlation between the latent factors and imposed the residual variance to be positive for Ezc.

**Supplementary Figure 2.**
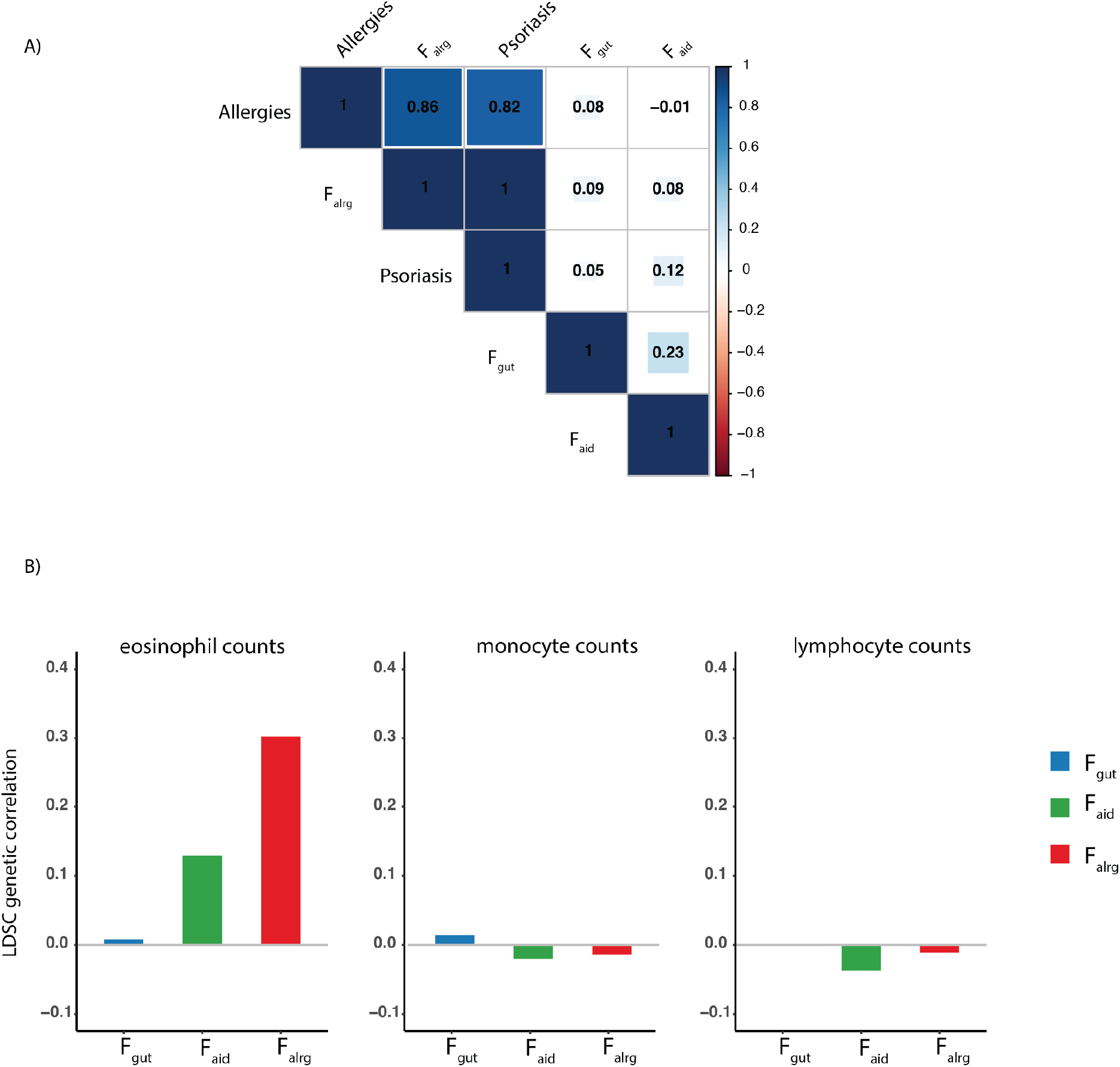
LDSC genetic correlations among Factors and allergic traits. **A)** LDSC genetic correlations among factors, psoriasis ^18^ and allergies ^19^. Shades of blue and red indicate positive and negative correlations respectively. **B)** LDSC genetic correlations between factors and circulating cell counts ^20^. Blue, green and red represent F_gut_, F_aid_ and F_alrg_ respectively.

**Supplementary Figure 3.**
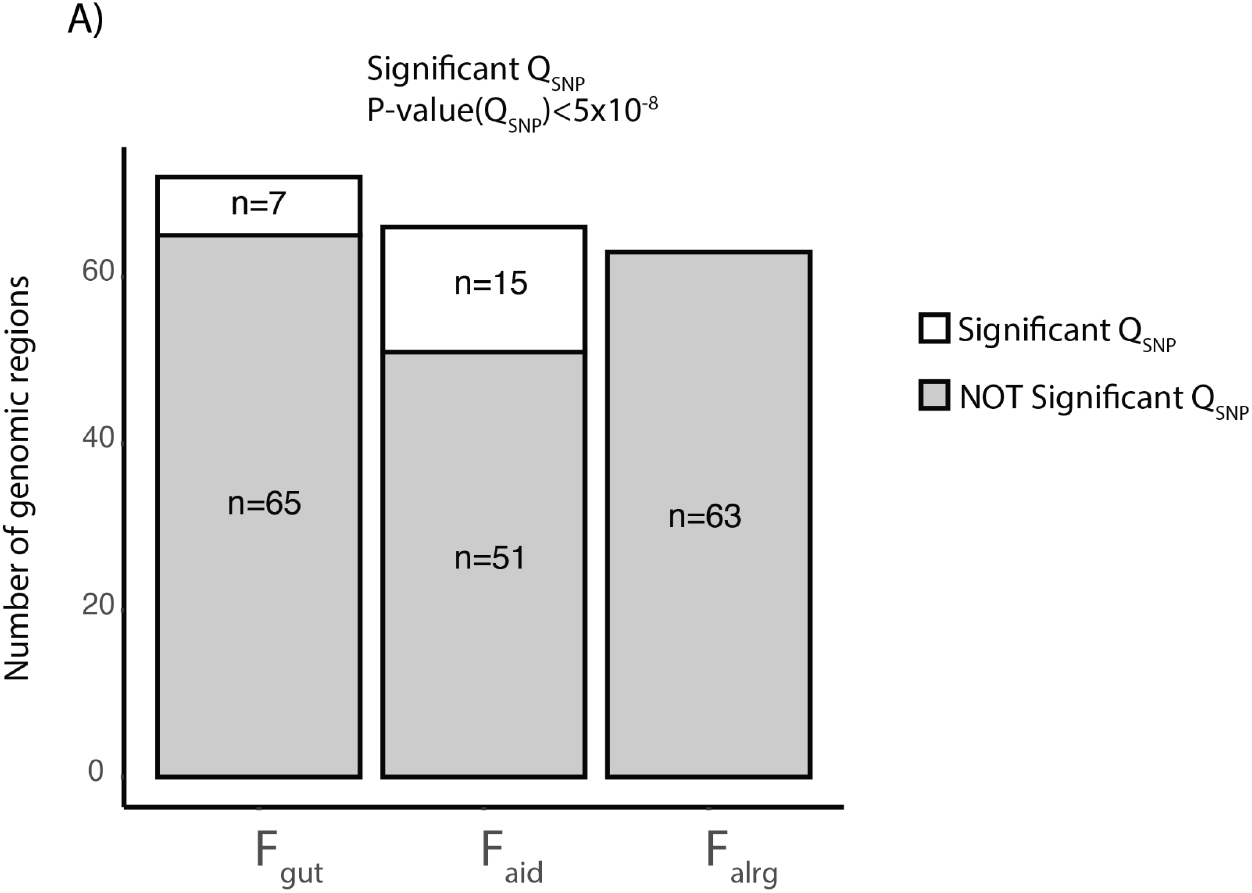
Qsnp statistics of genomic regions lead SNP. **A)** The bar plot shows the number of lead SNPs of the genomic region which had a significant Q_SNP_ (in white) and not significant (in grey).

**Supplementary Figure 4.**
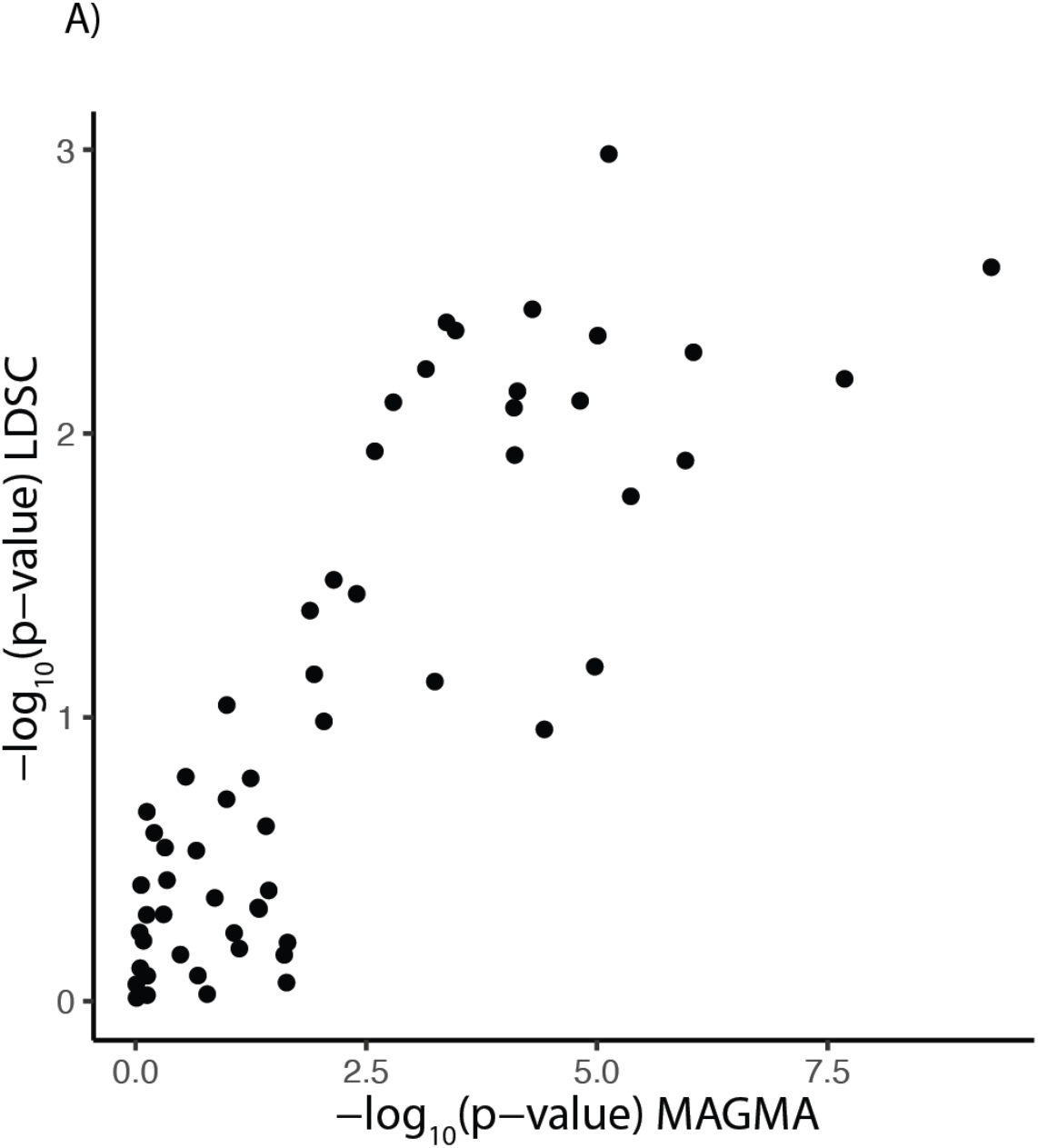
Comparison of LDSC and MAGMA enrichments. Dot plot shows correlation of −log_10_(p-value) between MAGMA and LDSC outputs for OneK1K cohort.

**Supplementary Figure 5.**
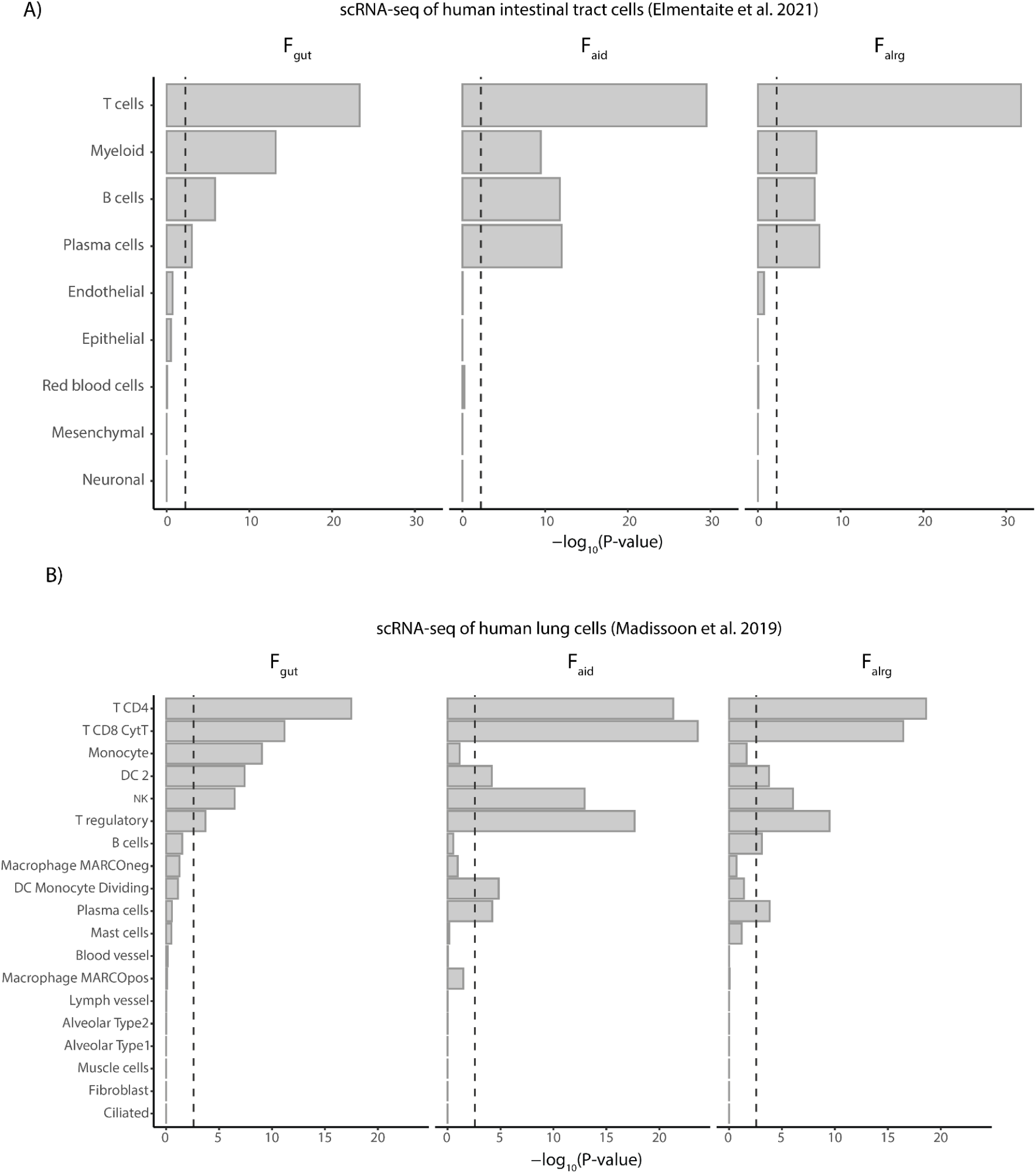
**A-B**) MAGMA gene-property results of intestinal cells ^28^ (**A**) and lung cells ^29^(**B**). The barplot shows −log_10_(p-value) of the enrichment.

